# The conformational equilibria of a human GPCR compared between lipid vesicles and aqueous solutions by integrative ^19^F-NMR

**DOI:** 10.1101/2024.10.14.618237

**Authors:** Arka Prabha Ray, Beining Jin, Matthew T. Eddy

**Author notes:** These authors have contributed equally to this work (Arka Prabha Ray, Beining Jin).

## Abstract

Endogenous phospholipids influence the conformational equilibria of G protein-coupled receptors, regulating their ability to bind drugs and form signaling complexes. However, most studies of GPCR-lipid interactions have been carried out in mixed micelles or lipid nanodiscs. Though useful, these membrane mimetics do not fully replicate the physical properties of native cellular membranes associated with large assemblies of lipids. We investigated the conformational equilibria of the human A_2A_ adenosine receptor (A_2A_AR) in phospholipid vesicles using ^19^F solid-state magic angle spinning NMR (SSNMR). By applying an optimized sample preparation workflow and experimental conditions, we were able to obtain ^19^F-SSNMR spectra for both antagonist- and agonist-bound complexes with sensitivity and linewidths closely comparable to those achieved using solution NMR. This facilitated a direct comparison of the A_2A_AR conformational equilibria across detergent micelle, lipid nanodisc, and lipid vesicle preparations. While antagonist-bound A_2A_AR showed a similar conformational equilibria across all membrane and membrane mimetic systems, the conformational equilibria of agonist-bound A_2A_AR exhibited differences among different environments. This suggests that the conformational equilibria of GPCRs may be influenced not only by specific receptor-lipid interactions but also by the membrane properties found in larger lipid assemblies.

## INTRODUCTION

Phospholipids are important endogenous allosteric modulators of G protein-coupled receptors (GPCRs), sensory integral membrane proteins essential for most physiological processes. As key components of biological membranes, phospholipids, along with cholesterol, have been frequently observed in close association with GPRCs in cryo-electron microscopy structures of GPCR signaling complexes^1–6^. These structural observations have been bolstered by biophysical approaches providing insights into the functional importance of GPCR-lipid interactions^7–11^. NMR spectroscopic studies, in particular, have shown that lipids not only associate with GPCRs but influence the conformational equilibria underpinning GPCR function, thereby modulating receptor activity in a manner that is highly dependent on the membrane environment^12–17^.

GPCR structures determined by cryo-EM^18,19^, as well as many biophysical experiments studying GPCR-lipid interactions^20–22^, have been carried out using either mixed micelles containing detergents and lipids or within lipid nanodiscs, membrane mimetics formed by a scaffold protein surrounding clusters of lipids^23,24^. Although nanodiscs have proven to be valuable biochemical tools, offering a more native-like alternative to detergents for solubilizing GPCRs, they still fall short of fully replicating the properties of biological membranes present in the cellular environment^25^. For example, the ability to alter membrane curvature, recognized for its role in regulating GPCR sorting^26^, is highly restricted in nanodiscs. Additionally, comparisons of lipid phase behavior have observed differences in the phase transitions of lipids between nanodiscs or membranes^27–31^, indicating that properties observed for larger assemblies of lipids are not replicated in nanodiscs. A comparison of cryo-EM structures of an ion channel among different sized nanodiscs revealed that the size of the lipid nanodisc influenced the determined structure^32^.

As an alternative to lipid nanodiscs, phospholipid vesicles exhibit properties such as coordinated lipid phase behavior and lipid composition that can be tailored to more accurately represent the same collective lipid properties found in native cellular membranes^33^. We leveraged this strength to investigate the conformational equilibria of the human A_2A_ adenosine receptor (A_2A_AR), a representative class A GPCR that has served as an important tool for GPCR biophysical studies^34–37^, in lipid vesicles using ^19^F magic angle spinning (MAS) solid-state NMR spectroscopy. ^19^F-NMR offers a unique advantage in solid-state MAS experiments due to its high sensitivity and minimal to no non-specific background signals, as documented in applications with integral membrane proteins^38,39^, membrane-associated proteins^40–43^ and protein assemblies^44,45^. To facilitate this investigation, we developed an optimized workflow for producing lipid vesicles containing A_2A_AR for ^19^F MAS NMR experiments. We confirmed that A_2A_AR in lipid vesicles maintained the same ligand binding affinities for antagonists and agonists as A_2A_AR in mammalian cells, and we confirmed that A_2A_AR was globally folded in our vesicle preparations. A distinct advantage of A_2A_AR prepared in lipid vesicles was enhanced thermal stability over A_2A_AR in either detergent micelles or lipid nanodiscs. By applying optimal sample preparation and experimental NMR conditions, we recorded ^19^F MAS SSNMR spectra of A_2A_AR with sensitivity and linewidths closely comparable to those observed for A_2A_AR in aqueous solutions. This enabled a direct comparison of the A_2A_AR conformational equilibria across lipid vesicles, lipid nanodiscs, and detergent micelle environments. While antagonist-bound A_2A_AR shared similar conformational equilibria across different environments, we observed differences in the populations of different conformations for agonist-bound A_2A_AR, suggesting that the bulk properties of lipids can impact the conformational landscape of GPCRs.

## RESULTS

### Reconstitution and pharmacological characterization of functional human A_2A_AR in lipid vesicles

For all biochemical and NMR experiments, a variant of human A_2A_AR was utilized containing a single, solvent-accessible cysteine located at position 289 at the intracellular surface of transmembrane (TM) helix VII, A_2A_AR[A289C]. Earlier NMR studies demonstrated that spectra of A_2A_AR[A289C] labeled with ^19^F-2,2,2-trifluoroethanethiol at C289, A_2A_AR[A289C^TET^] were sensitive to differences in the efficacies of bound ligands^46^ and sensitive to changes in the lipid composition within phospholipid nanodiscs^14–16^. The chemical shift of the ^19^F-NMR probe located at position 289 was shown to be sensitive to ring current effects from nearby aromatic residues F286^7.51^,F295^8.50^, and F299^8.54^ (superscripts denote the Ballosteros-Weinstein nomenclature^47^), which appear near the interface between transmembrane helix VII and the amphipathic helix VIII. These residues have been found to be conserved among other GPCRs and have been proposed to form hydrophobic interactions with nearby hydrophobic residues in TM I, forming a local microswitch important to G protein signaling^48^. Thus, the ^19^F NMR probe located at position 289 reports on structural changes linked to global rearrangements related to GPCR activation. The same position and labeling protocol utilized in earlier studies was applied to the current study to facilitate comparison of the A_2A_AR[A289C^TET^] conformational equilibria between vesicles, lipid nanodiscs, and detergent environments.

We developed an optimal workflow to prepare phospholipid vesicles containing A_2A_AR[A289C] for ^19^F MAS NMR experiments (Figure 1). The workflow followed from earlier studies that reconstituted detergent-solubilized transmembrane proteins into lipid vesicles^49–51^. The general approach was to first prepare vesicles of defined phospholipid composition and size, then destabilize the vesicles using detergent to facilitate insertion of purified, detergent-solubilized A_2A_AR[A289C], selectively remove the detergent, and homogenize the vesicles one final time before biochemical or NMR experiments (Figure 1). Phospholipid vesicles with and without reconstituted A_2A_AR[A289C] were characterized as described in the following text.

**Figure 1.**
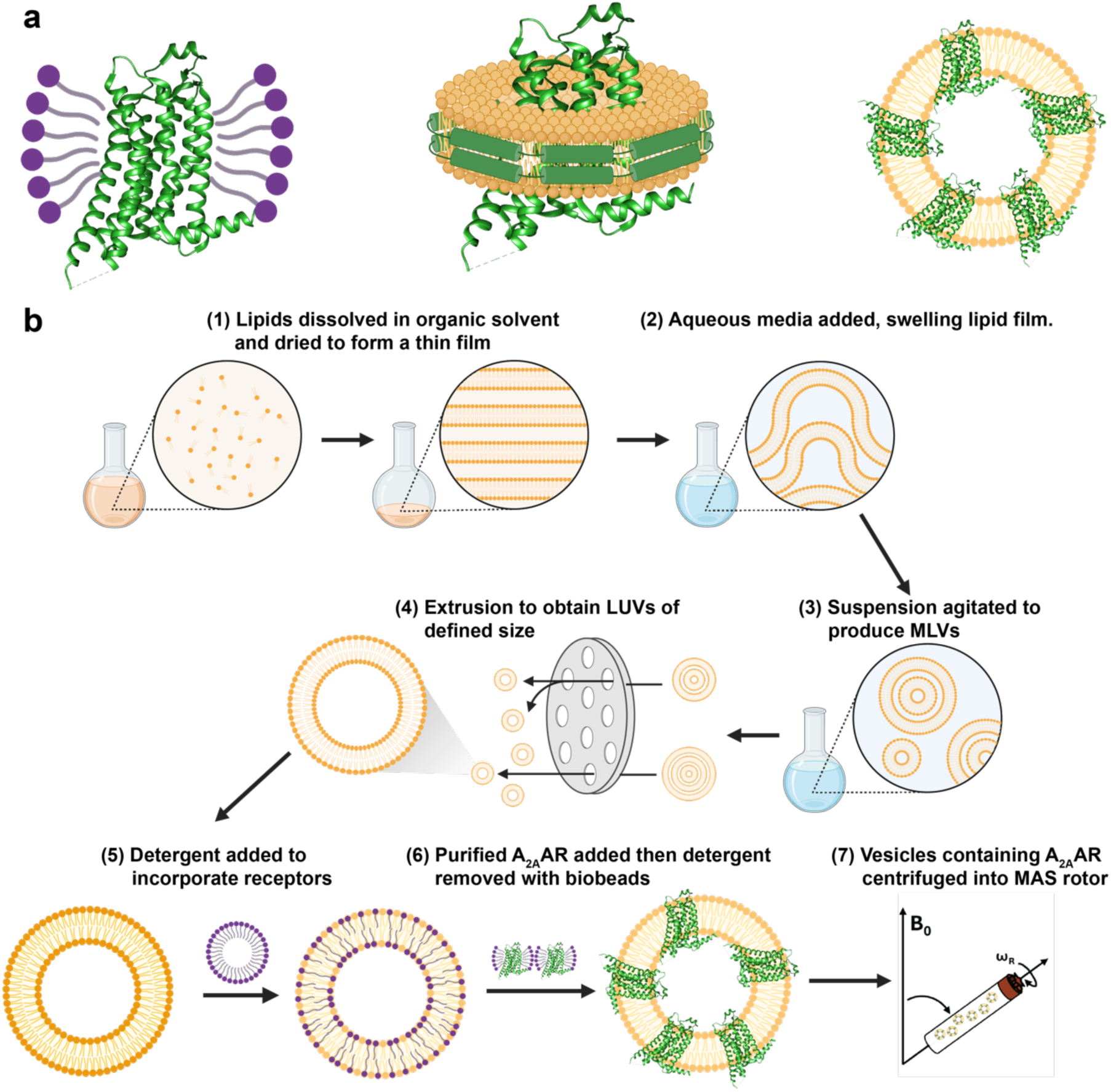
Human A_2A_AR in different membrane or membrane-mimetic environments and sample preparation workflow. (a) A_2A_AR compared in three different environments in this study: (left) detergent micelles, (middle) lipid nanodiscs, and (right) lipid vesicles. (b) Schematic of the sample preparation workflow for preparing lipid vesicles containing ^19^F-labeled human A_2A_AR for solid-state MAS NMR experiments.

To verify formation of phospholipid vesicles, we recorded negative stain electron micrograph images of vesicles without and with reconstituted A_2A_AR[A289C] (Figure 2a and 2b). Vesicle size was confirmed to be consistent with our expectations of ∼200 nm, based on the pore size of 200 nm used for extrusion of the vesicles. Average vesicle diameters and distribution of vesicle sizes were confirmed in dynamic light scattering (DLS) experiments. Vesicles with and without reconstituted A_2A_AR were homogenous in size, with a polydispersity index of < 0.1 for vesicles without A_2A_AR and a polydispersity index < 0.2 for vesicles containing A_2A_AR (Figure 2c).

**Figure 2.**
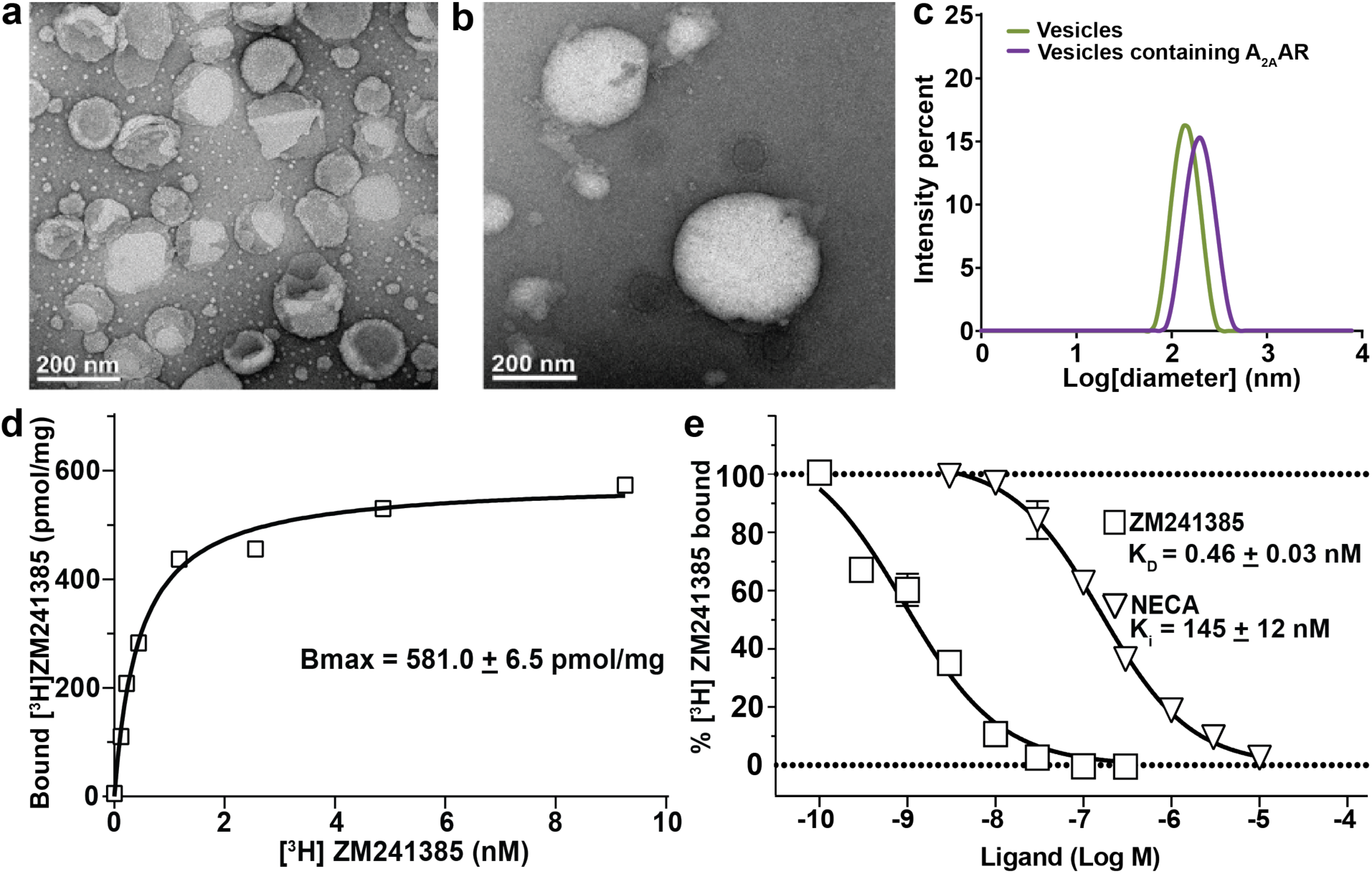
Characterization of lipid vesicles containing human A_2A_AR and pharmacological validation of A_2A_AR in vesicles. (**a and b**) Representative negative stain electron micrographs of unilameller vesicles composed of POPC and POPS (70:30 molar ratio) (**a**) without and (**b**) with reconstituted A_2A_AR. (**c**) Dynamic light scattering measurements of the distribution of the sizes of vesicles composed of POPC and POPS (70:30 molar ratio) without (green) and with (purple) A_2A_AR. (**d and e**) Pharmacological characterization of A_2A_AR in lipid vesicles. (**d**) Saturation binding experiment with ^3^H-ZM241385 and A_2A_AR in lipid vesicles containing POPC and POPS (70:30 molar ratio). The reported B_max_ value represents the mean and associated error is the s.e.m. from 3 independent trials done in triplicate. (**e**) Radioligand competition experiments. K_D_ and K_i_ values are reported for the antagonist ZM241385 and the agonist NECA, respectively. The reported error represents the s.e.m. from 3 independent trials done in triplicate.

To confirm the pharmacological function of A_2A_AR[A289C] in lipid vesicles, we recorded radioligand saturation binding experiments and radioligand competition binding experiments (Figure 2d and 2e). In saturation binding experiments, increasing concentrations of [^3^H]ZM241385 were incubated with POPC/POPS (70:30 molar ratio) vesicles containing A_2A_AR[A289C], and specific binding was determined as the difference in observed binding between samples prepared in the absence and presence of 10 µM cold ZM241385. From this method, we determined a B_MAX_ value for A_2A_AR[A289C] in lipid vesicles to be 581.0 ± 6.5 pmol/mg (Figure 2d). This value is approximately a 22-fold increase in the amount of functional receptors as compared with A_2A_AR expressed in the plasma membranes of insect cells^52^, confirming that our vesicle samples contained fully functional receptors. For competition binding experiments, vesicle preparations were incubated with [^3^H]-ZM241385 and increasing concentrations of cold ZM241385 or NECA. From competition binding experiments, we determined the affinities of A_2A_AR[A289C] in POPC/POPS vesicles for a representative antagonist and agonist, determining a K_D_ of 0.46 nM for the antagonist ZM241385 and K_I_ for the agonist NECA of 145 nM (Figure 2e), nearly identical to the affinities determined for A_2A_AR in insect cells^52^.

### Anionic lipids induce asymmetric insertion of A_2A_AR into lipid vesicles

Earlier structural characterization of membrane proteins in lipid vesicles have shown that integral membrane proteins can adopt multiple global orientations within vesicles^53^. Reconstituted A_2A_AR could potentially adopt two different global orientations: an “outward facing” orientation, where the orthosteric binding pocket facing away from the vesicle interior, or an “inward facing” orientation, where the orthosteric binding pocket is directed toward the vesicle interior (Figure 3). To determine the orientation of A_2A_AR[A289C] reconstituted in lipid vesicles, we adapted a previously reported protocol for quantifying the orientation of integral membrane proteins within phospholipid vesicles^54^. Following this method, a cyanine fluorophore was covalently attached to a single solvent accessible cysteine C289, the same position used for ^19^F-TET labeling in NMR experiments, on A_2A_AR purified in detergent. The labeled receptor was then reconstituted into vesicles, and the fluorescence intensity was measured. Next a membrane-impermeable fluorescence quencher was introduced, and the resulting decrease of fluorescence reported on receptors oriented with the fluorescent dye facing outward, away from the vesicle. Finally, detergent is added to disrupt the vesicles, and fluorescence is measured again to quantify the receptors with an inward facing orientation (Figure 3).

**Figure 3.**
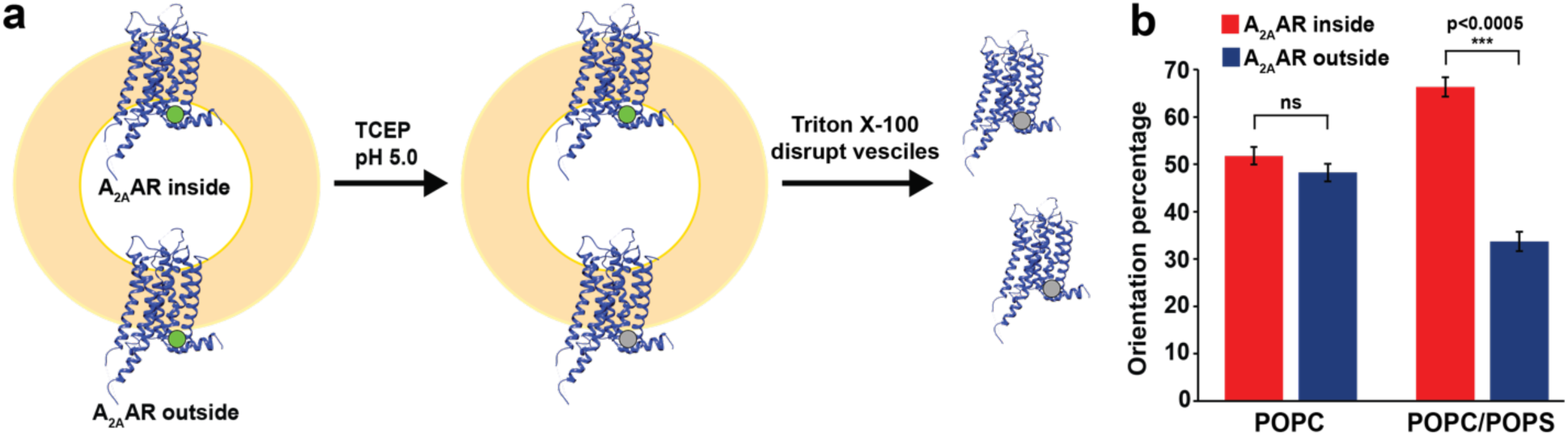
Determination of the orientation of A_2A_AR in lipid vesicles. (**a**) Schematic of the fluorescence-quenching assay used to quantify receptor orientation within vesicles. Green circles represent position C289 labeled with Cy3. Receptors with Cy3 covalently attached to position C289 facing toward the vesicle interior are labeled “A_2A_AR inside”, while receptors with Cy3 facing away from the vesicle interior are labeled “A_2A_AR outside”. Grey circles represent Cy3-labels that have been chemically quenched. (**b**) Quantitative comparison of the orientation of A_2A_AR in vesicles made from POPC or POPC and POPS (70:30 molar ratio). The orientations “A_2A_AR inside” and “A_2A_AR outside” are as defined in (**a**). Error bars indicate the s.e.m. calculated from n≥3 independent experiments. Statistically significant values are illustrated as ***p<0.005 using a 2-tailed unpaired t-test.

This approach was applied to quantifying the orientation of A_2A_AR[A289C] in vesicles containing two different lipid compositions, vesicles composed of POPC and vesicles composed of a defined binary mixture of POPC and POPS at a 70:30 molar ratio. For A_2A_AR[A289C] reconstituted into vesicles containing POPC, the orientation distribution was found to show a nearly equal distribution of outward facing and inward facing receptors (Figure 3). In contrast to this, for A_2A_AR[A289C] reconstituted into vesicles composed of POPC and POPS (70:30 molar ratio), approximately 70% of the receptors adopted an outward facing orientation while approximately 30% of receptors adopted an inward facing orientation (Figure 3).

Because the POPC and POPC/POPS vesicles were prepared using identical protocols that created homogenous vesicles of closely similar overall size, the differences observed in receptor orientation are unlikely to be due to variation in vesicle size. Instead, the receptor preference for an outward facing orientation in vesicles composed of a binary mixture of POPC and POPS is more attributed to differences in membrane properties, such as lipid order^55,56^ or local curvature induced by addition of POPS^57^. Additionally, the asymmetry of receptor orientation in POPC/POPS vesicles suggests the radioligand saturation data shown in Figure 2 may have undercounted the number of functional receptors. This is likely because the ^3^H-labeled antagonist ZM241385, like most other A_2A_AR ligands, may not fully penetrate the vesicles. Therefore, the estimated number of functional receptors in our NMR samples ranges from approximately 580 to 750 pmol/mg.

### Human A_2A_AR is more thermally stable in lipid vesicles than in nanodiscs or detergent micelles

To verify that A_2A_AR[A289C] reconstituted into lipid nanodiscs was globally folded, we recorded microscale fluorescence thermal melting assays by adapting a previously reported protocol for detergent-solubilized GPCRs.^58^ A_2A_AR[A289C] in complex with the antagonist ZM241385 reconstituted in vesicles composed of POPC and POPS (70:30 molar ratio) exhibited a thermal melting curve that showed a sharp transition from folded to unfolded protein, indicating cooperative unfolding consistent with a folded receptor sample population (Figure 4). From these experiments, the melting temperature (T_M_) of A_2A_AR[A289C] in complex with the antagonist ZM241385 in lipid vesicles composed of POPC and POPS (70:30 molar ratio) was determined to be 77.8 ± 0.5 °C. This was found to be significantly higher than the melting temperatures of A_2A_AR[A289C]in complex with ZM241385 in either lipid nanodiscs composed of the same ratio of POPC and POPS (T_M_ 68.6 ± 0.5 °C) or in DDM/CHS mixed micelles (T_M_ 61.2 ± 0.7 °C). Thus, the lipid vesicle environment provided the highest possible thermal stability for A_2A_AR among all membrane or membrane mimetic systems.

**Figure 4.**
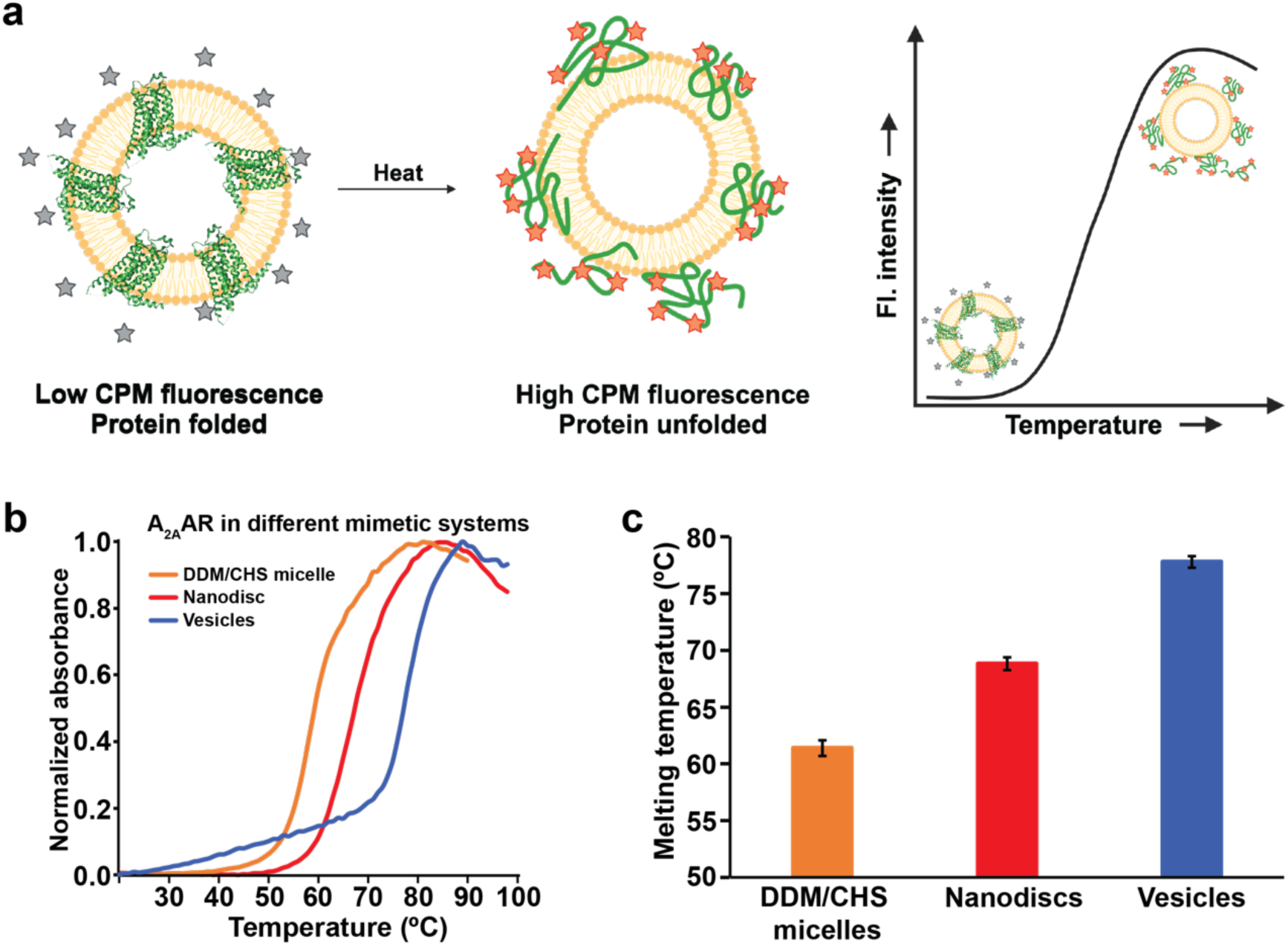
Fluorescence thermal melting profiles of the A_2A_AR complex with the antagonist in three different membrane mimetics. (**a**) Schematic of the fluorescence thermal shift assay as applied to A_2A_AR in lipid vesicles. The inactive fluorescent dye (grey stars) shows increased emission upon covalent attachment with cysteines that become solvent accessible upon protein unfolding (orange stars). (**b**) Representative thermal melting profiles for A_2A_AR in DDM/CHS detergent micelles, lipid nanodics containing POPC and POPS (70:30 molar ratio), and lipid vesicles containing POPC and POPS (70:30 molar ratio). (**c**) The melting temperature for A_2A_AR in each membrane mimetic is reported as the mean of three independent experiments ± s.e.m.

### The conformational equilibrium of antagonist-bound A_2A_AR in lipid vesicles

We observed the conformational equilibria of A_2A_AR[A289C^TET^] in complex with the antagonist ZM241385 in lipid vesicles composed of a defined binary mixture of POPC and POPS (70:30 molar ratio) utilizing the protocol from Figure 1. This lipid composition was selected to facilitate comparisons between the present solid-state NMR studies of A_2A_AR[A289C^TET^] in lipid vesicles with earlier solution NMR studies of A_2A_AR[A289C^TET^] in detergent micelles^46^ and lipid nanodiscs containing the same ratio of POPC and POPS^14–16^.

The ^19^F-MAS NMR spectra of A_2A_AR[A289C^TET^] recorded at 20 kHz MAS with 105 kHz ^1^H TPPM decoupling contained two signals at δ ≈ 11.3 ppm (P3) and δ ≈ 9.5 ppm with the signal P3 being the largest signal in the spectrum (Figure 5a and b, Figure S1). The shape of the overall ^19^F signal envelop, the number of observed spectral components, the chemical shifts of the two observed components, and the relative intensities of the populations P1 and P3 appeared highly similar between the solid-state preparation of antagonist-bound A_2A_AR[A289C^TET^] in lipid vesicles and antagonist-bound A_2A_AR[A289C^TET^] in lipid nanodiscs or detergent micelles in aqueous solutions (see below). With 105 kHz ^1^H TPPM decoupling, we observed marginal decreases in the line widths for both components with increasing MAS frequency (Figure 5a, 5b and 5g), with a marginal increase in the linewidth for component P1 at 40 kHz MAS.

**Figure 5.**
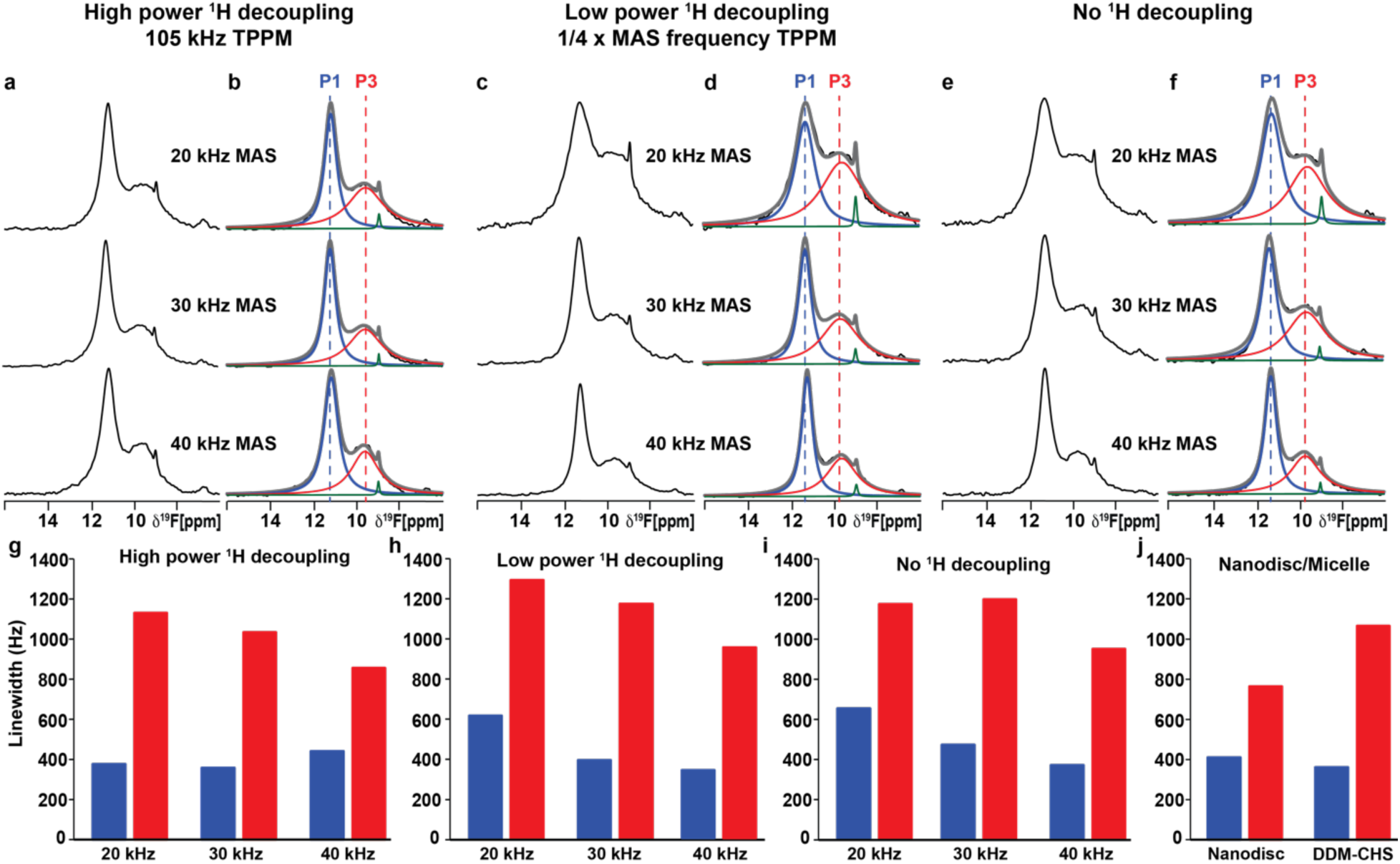
^19^F MAS NMR-observed conformational equilibria of human A_2A_AR in complex with an antagonist in lipid vesicles measured with different experimental parameters. (**a**) 1-dimensional ^19^F-MAS spectra of A_2A_AR[A289C^TET^] in complex with the antagonist ZM241385 reconstituted into POPC/POPS lipid vesicles recorded with 105 kHz ^1^H TPPM decoupling at three different MAS frequencies. (**b**) NMR spectra from (a) are shown superimposed with Lorentzian deconvolutions with the minimal number of components that provided a good fit, labeled P1 and P3. (**c**) 1-dimensional ^19^F-MAS spectra of the same sample used to measure the data in (a) recorded with ^1^H TPPM decoupling power set to one-quarter of the applied MAS frequency. (**d**) NMR spectra from (c) are shown superimposed with Lorentzian deconvolutions with the minimal number of components that provided a good fit, labeled P1 and P3. (**e and f**) (**e**) 1-dimensional ^19^F-MAS spectra of the same sample used to measure the data in (a) recorded with no ^1^H decoupling and (**f**) NMR spectra from (**e**) are shown superimposed with Lorentzian deconvolutions. (**g-j**) Linewidths measured for populations P1 and P3 for A_2A_AR in (**g-i**) lipid vesicles or (**j**) lipid nanodiscs or detergent micelles. Components colored green are from free TET, consistent with earlier NMR studies (see text).

Previously reported SSNMR studies of microcrystalline amino acids and proteins demonstrated that at moderate to higher MAS frequencies, the application of low power TPPM ^1^H decoupling with an applied field of one-quarter of the MAS frequency was effective at providing high resolution spectra in ^13^C-detected experiments^59,60^. We tested whether ^1^H TPPM decoupling with an applied field of one-quarter of the MAS frequency would produce spectra with resolution similar to that obtained with high power TPPM decoupling. At 20 kHz MAS frequency, the lines in ^19^F-NMR spectra recorded with low power ^1^H TPPM were about 20% broader than what we observed at the same spinning frequency with high power ^1^H TPPM (Figure 5c, 5d and 5h). With increasing spinning frequency, we observed narrowing of both spectral components with low power ^1^H TPPM decoupling, from ∼1300 Hz for P3 and ∼620 Hz for P1 at 20 kHz MAS to ∼940 Hz for P3 and ∼340 Hz for P1 at 40 kHz MAS frequency (Figure 5c, 5d, 5h and Table S2). As a point of comparison, we also recorded ^19^F MAS SSNMR spectra without any applied ^1^H decoupling over the same range of MAS frequencies (Figure 5e and 5f). At 20 kHz MAS frequency, we observed broader lines without decoupling, especially for component P3, which had a measured line width of ∼650 Hz (Table S2). At 40 kHz MAS frequency, we observed significant narrowing for both components, ∼370 Hz for component P3 and ∼950 Hz for component P1 (Table S2), which was only marginally broader than the line width for P1 observed with low power ^1^H decoupling and high power ^1^H decoupling (Table S2). Considering the line widths for both components P1 and P3, we observed the overall best spectra resolution at 40 kHz MAS with low power ^1^H TPPM decoupling (Figure 5c and 5d).

We then used the optimal experimental conditions, 40 kHz MAS frequency and low power ^1^H TPPM decoupling, to compare the conformational equilibria of antagonist-bound A_2A_AR[A289C^TET^] across a range of sample temperatures (Figure S2). ^19^F MAS SSNMR spectra were recorded for antagonist-bound A_2A_AR[A289C^TET^] in lipid vesicles containing POPC and POPS (70:30 molar ratio) at temperature set points ranging from 245 K to 275 K. These temperatures represent the temperatures the cooling unit was set to and not the actual sample temperature. To obtain a more accurate measurement of the sample temperature, we accounted for the frictional heating caused by magic angle spinning, following established protocols that use the chemical shift of ^79^Br in KBr powder^61^. At 40 kHz MAS, frictional heating was found to increase the sample temperature by approximately 40 °C. Therefore, the actual sample temperatures investigated were closer to 285 K and 315 K. Because low power ^1^H TPPM deposits significantly less radiofrequency energy into the sample, the estimated sample temperatures accounting for frictional heating are likely very close to the actual sample temperatures. Across this temperature range, we observed only minor differences in the line widths or relative populations of the two components in all spectra of antagonist-bound A_2A_AR[A289C^TET^] (Figure S2).

### The conformational equilibrium of agonist-bound A_2A_AR in lipid vesicles

To investigate the conformational equilibria of agonist-bound A_2A_AR[A289C^TET^] in lipid vesicles, we prepared a sample in vesicles with the same defined binary composition (POPC and POPS, 70:30 molar ratio) in complex with the full agonist NECA, a high-affinity adenosine derivative. Given that we obtained the best overall spectral resolution using low power ^1^H TPPM decoupling for antagonist-bound A_2A_AR[A289C^TET^], we applied the same conditions to record ^19^F-SSNMR spectra of NECA-bound A_2A_AR[A289C^TET^] at three different MAS frequencies (Figure 6).

**Figure 6.**
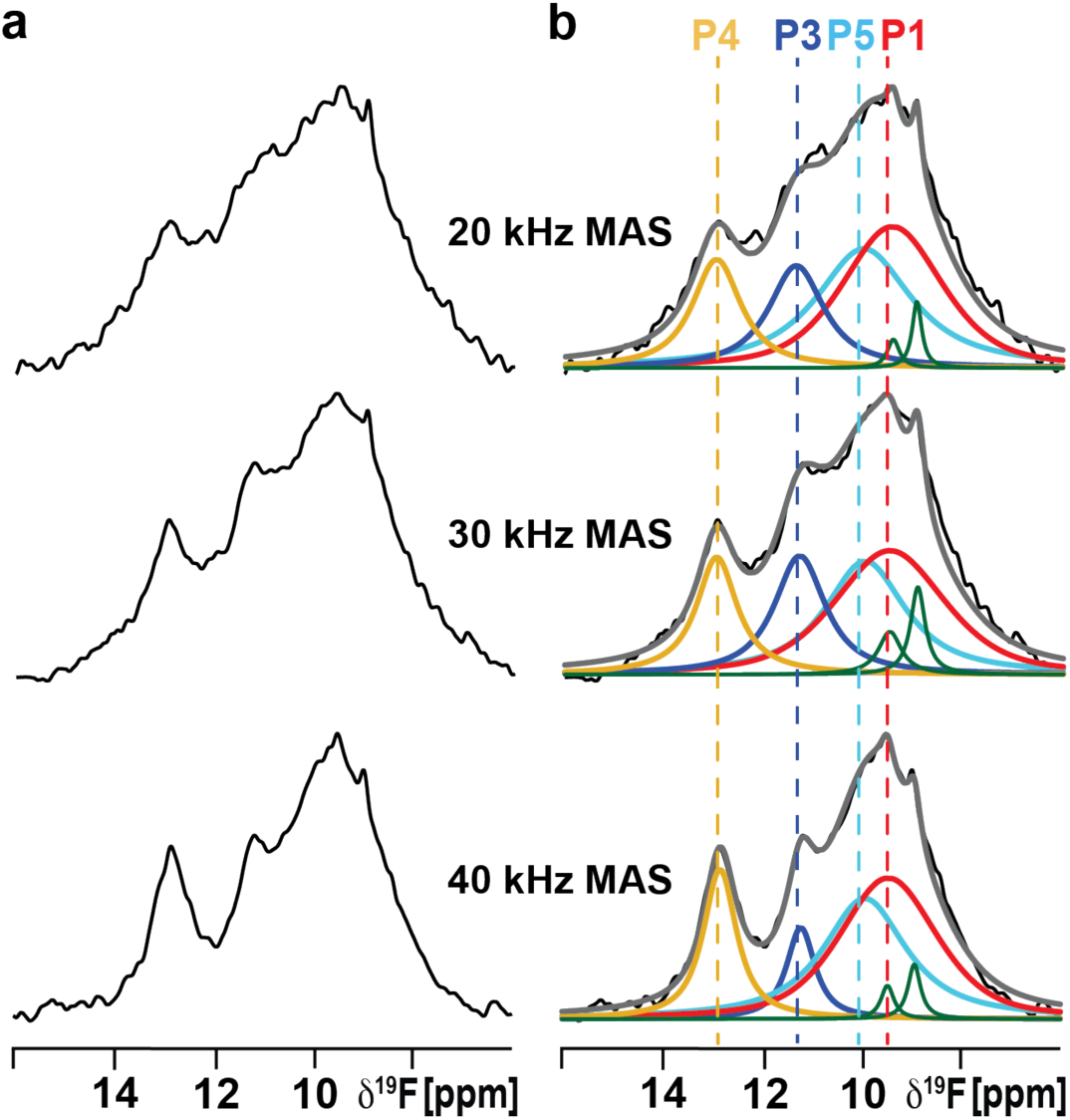
^19^F MAS NMR-observed conformational equilibria of human A_2A_AR in complex with the agonist NECA in lipid vesicles measured with low power TPPM ^1^H decoupling at several MAS frequencies. (**a**) 1-dimensional ^19^F-MAS spectra of A_2A_AR[A289C^TET^] in complex with the agonist NECA reconstituted into POPC/POPS lipid vesicles recorded with three different MAS frequencies and ^1^H TPPM decoupling at an applied ^1^H power of one-quarter of the MAS frequency. (**b**) NMR spectra from (a) are shown superimposed with Lorentzian deconvolutions with the minimal number of components that provided a good fit, labeled P1 through P5. Components colored green are from free TET, consistent with earlier NMR studies (see text).

At 20 kHz MAS frequency, the ^19^F-MAS NMR spectra of A_2A_AR[A289C^TET^] in complex with the agonist NECA contained multiple components with somewhat overlapped broader lines (Figure 6). Increasing the MAS frequency significantly improved the resolution, narrowing the line widths of all components at 30 kHz MAS and further at 40 kHz MAS (Figure 6). Between 20 kHz and 40 kHz MAS, the line widths of most components were narrowed by 30% to 50%. At 40 kHz MAS, we were able to record ^19^F MAS SSNMR spectra with distinct spectral components (Figure 6) that facilitated deconvolution of the spectrum. The chemical shifts of the individual components were almost identical between the solid-state preparation of agonist-bound A_2A_AR[A289C^TET^] in lipid vesicles and agonist-bound A_2A_AR[A289C^TET^] in lipid nanodiscs or detergent micelles in aqueous solutions. Interestingly the relative populations of the individual components in SSNMR spectra of vesicle preparations differed from those observed in solution, as discussed further below.

### Comparing the conformational equilibrium of A_2A_AR across three different membrane or membrane-mimetic systems

The sensitivity and resolution obtained in ^19^F MAS SSNMR spectra of lipid vesicle preparations of A_2A_AR[A289C^TET^] complexes with both the antagonist ZM241385 and the agonist NECA measured at 40 kHz MAS approached the resolution observed in spectra of A_2A_AR[A289C^TET^] detergent or nanodisc preparations in aqueous solutions (Figure 7). The similarity of the chemical shifts for all spectra components facilitated a direct comparison of the conformational equilibria of A_2A_AR across the three different membrane or membrane-mimetic systems (Figure 7).

**Figure 7.**
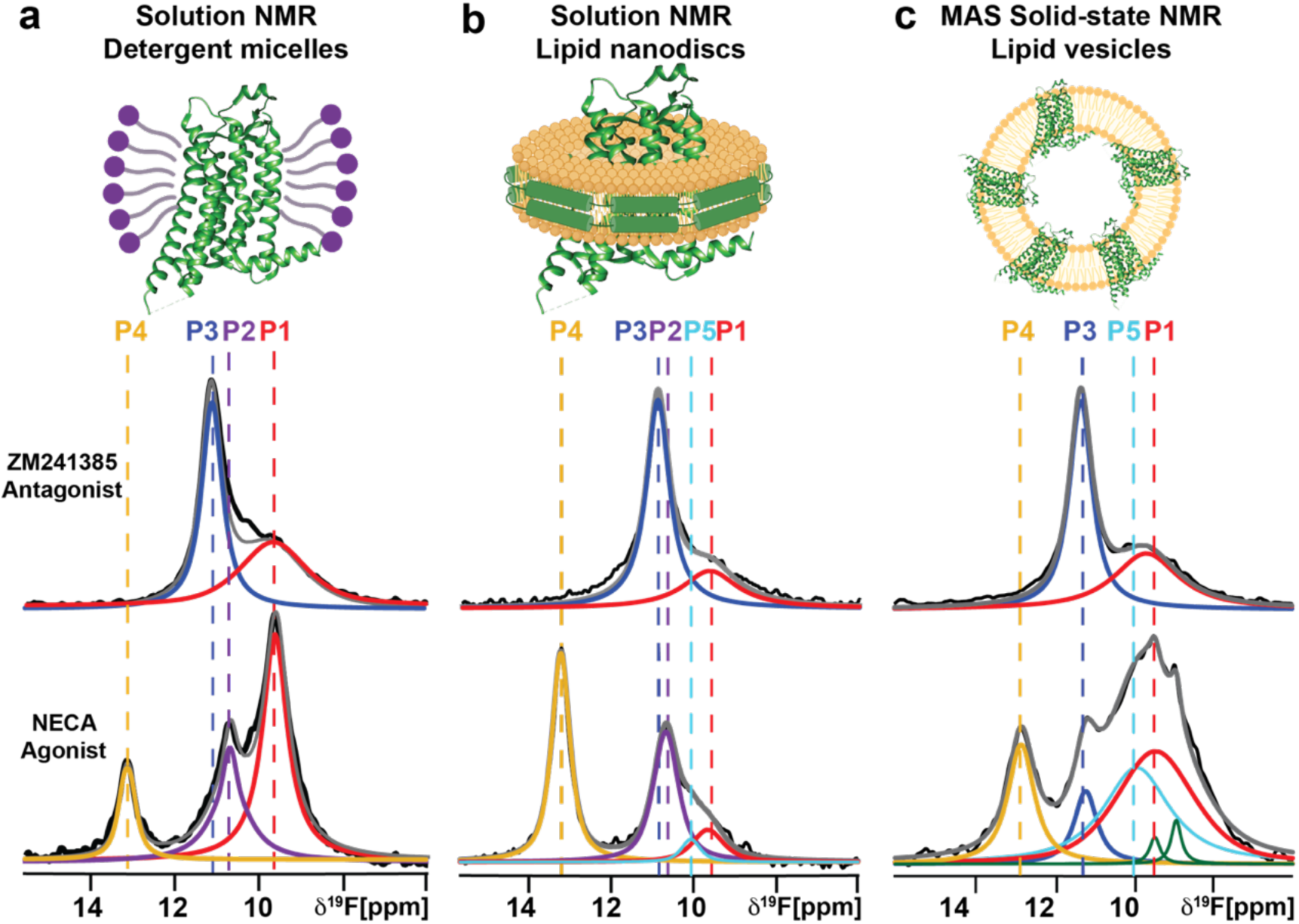
^19^F-NMR systematic comparison of the conformational equilibria of antagonist-bound and agonist-bound human A_2A_AR[A289C^TET^] across three membrane or membrane-mimetic systems by solution NMR in **(a)** detergent micelles and **(b)** lipid nanodiscs and by MAS solid-state NMR in **(c)** lipid vesicles. Same color scheme as in Figures 5 and 6.

For antagonist-bound A_2A_AR[A289C^TET^], we observed highly similar spectra for lipid vesicle, lipid nanodisc, and detergent micelle preparations. In all spectra, we observed two components, P1 and P3, that exhibited nearly identical chemical shifts. The relative intensities of both components, and thus the relative population of each conformation, varied only marginally across all sample conditions (Figure 7). This indicated that the conformational equilibria of antagonist-bound A_2A_AR was largely consistent between samples prepared in lipid vesicles and lipid nanodiscs of the same binary lipid composition and detergent micelles.

In contrast, for agonist-bound A_2A_AR[A289C^TET^], we observed differences in the number of components in each spectrum among the different sample preparation conditions. The chemical shifts of the components remained nearly identical across all samples. Additionally, the relative populations of each component varied between lipid vesicles and the two membrane mimetic systems (Figure 7). This indicated that the conformational equilibrium of agonist-bound A_2A_AR was much more sensitive to changes in the membrane environment than antagonist-bound A_2A_AR.

## DISCUSSION

The data presented in Figures 2 and 4 demonstrate advantages of structural and biophysical investigations of a human receptor proteins in lipid vesicles, specifically the ability to generate homogenous samples of pharmacologically active A_2A_AR that are more thermally stable than preparations in detergent micelles or lipid nanodiscs. GPCRs, and membrane proteins generally, are often difficult to work with because of their relatively lower thermal stability, prompting the development of alternative detergents and other membrane mimetics to facilitate biophysical studies^62^. Results from Figure 4 show that vesicles provide an improved level of thermal stability for A_2A_AR, and the benefits of vesicle preparations in providing a stabilizing environment can likely be extended to additional GPCRs.

Applying an optimized protocol to prepare samples of human A_2A_AR in lipid vesicles, we were able to record ^19^F-NMR solid-state MAS spectra with resolution and sensitivity closely comparable to that obtained in solution with A_2A_AR in detergent micelles or lipid nanodiscs (Figures 5-7). This means we can record spectra with quality comparable to what we can obtain in solution, with the significant advantage that we can more closely replicate properties of the cellular membrane in lipid vesicles. In optimal experimental conditions for spectra of antagonist-bound A_2A_AR[A289C^TET^], measured with low power TPPM ^1^H decoupling and 40 kHz MAS frequency, the line width for component P1 was only ∼15% wider than the line width for the same component measured in lipid nanodiscs, and the line width for component P3 was slightly narrower in MAS SSNMR spectra than measured in lipid nanodiscs (Figure 5). The narrowing of line widths for both components with increasing MAS without ^1^H decoupling suggested that the ^19^F chemical shift anisotropy (CSA) was likely a larger contributing factor than ^1^H-^19^F dipolar couplings. We anticipate that by applying even higher MAS frequencies, we may expect to see additional line narrowing, as has been demonstrated in ^19^F SSNMR experiments of model compounds recorded with MAS frequencies between 60 and 111 kHz^63^.

The high resolution achieved in our ^19^F SSNMR MAS experiments enabled a direct comparison of the conformational equilibrium of human A_2A_AR in solid-state lipid vesicle samples with that in aqueous solution preparations using detergents or lipid nanodiscs (Figure 7). This systematic comparison revealed both similarities and differences in the A_2A_AR conformational equilibria across the three membrane or membrane mimetic systems. For antagonist-bound A_2A_AR, the spectral envelop exhibited the same number of components with nearly identical chemical shifts in both solid-state lipid vesicle samples and in aqueous solution preparations with DDM/CHS detergent micelles and lipid nanodiscs (Figure 5). This indicated that the conformation of antagonist-bound A_2A_AR near the intracellular surface is highly similar across all three environments. In contrast, for agonist-bound A_2A_AR, we observed differences in the ^19^F-NMR signal envelop among the three different environments. Specifically, the relative populations of each conformer varied considerably (Figure 7). The ^19^F-NMR spectra of A_2A_AR[A289C^TET^] in complex with NECA measured in lipid vesicles and lipid nanodiscs employed the same defined binary lipid composition of POPC and POPS in the same ratio. Therefore, the observed differences in the relative populations of each conformation between lipid nanodisc and lipid vesicle preparations may reflect the influence of distinct bulk lipid properties on the conformational equilibria of A_2A_AR. This raises intriguing possibilities for further investigation, particularly in additional GPCR systems, where variations in lipid properties—such as membrane curvature—could be systematically explored using vesicles to assess their impact on receptor conformational equilibria.

## METHODS

### Molecular cloning

The gene encoding human A_2A_AR (1-316), cloned into a pPIC9K vector (Invitrogen) at the BamHI and NotI restriction sites, contained a single amino acid replacement (N154Q) to remove the only glycosylation site, an N-terminal FLAG tag, and a 10 X C-terminal His tag. PCR-based site-directed mutagenesis was used to replace A289^7.54^ with cysteine, creating A_2A_AR[A289C] using the Accuprime *Pfx* SuperMix (ThermoFisher Scientific). The construct design is consistent with earlier studies^15,16,46,64^, and no new constructs were generated for this study.

### A_2A_AR production

Plasmids containing A_2A_AR[A289C] were transformed into the BG12 strain of *Pichia pastoris* (Biogrammatics) via electroporation. High-expressing clones were selected via small-scale protein expression screening method where protein expression was evaluated by an anti-FLAG western blot assay, as previously reported^64,65^. Glycerol stocks of the high expressing clones were prepared and stored at -80 °C for future use.

A_2A_AR was expressed in *P. pastoris* following previously published protocols^15,64^. 4 mL cultures in buffered minimal glycerol (BMGY) media were inoculated from glycerol stocks and grown at 30 °C for 48 h. These cultures were used to inoculate 50 mL of BMGY medium and grown at 30 °C for an additional 60 h. Next, the cultures were then used to inoculate 500 mL of BMGY medium and grown for 48 h at 30 °C. The cells were then harvested by centrifugation and resuspended in 500 mL of buffered minimal methanol (BMMY) media without methanol. The temperature was lowered to 28 °C and no additional carbon source was added for 6 h to consume any remaining glycerol before protein expression was induced by addition of methanol to a final concentration of 0.5% (w/v). Two more aliquots of methanol were added at 12 h intervals after induction for a total expression time of 36 h. The cells were then harvested by ultracentrifugation at 3000 * g, and the cell pellets were frozen in liquid nitrogen and stored at -80 °C for future use.

### A_2A_AR purification and ^19^F-labeling via chemical modification

Purification and ^19^F-labeling via an in-membrane chemical modification (IMCM) approach was carried out following previously reported protocols^66^. Cell pellets were resuspended in lysis buffer (100 mM NaCl, 50 mM sodium phosphate pH 7.0, 5% glycerol (w/v), and in-house prepared protease inhibitor solution) and lysed using a cell disruptor (Pressure Biosciences) at 40k PSI. Cell membranes containing A_2A_AR[A289C] were isolated by ultracentrifugation at 200,000 * g for 30 min and homogenized in buffer (10 mM HEPES pH 7.0, 10 mM KCl, 20 mM MgCl_2_, 1 M NaCl, 4 mM theophylline). The homogenized membranes were incubated with 1 mM of 4,4’-dithiodipyridine (aldrithiol-4) and protease inhibitor cocktail solution (in-house prepared) for 1 h at 4 °C. Membrane suspensions were pelleted by ultracentrifugation at 200,000 * g for 30 min and the supernatant was discarded to remove excess aldrithiol-4. The pelleted membranes were resuspended in the same buffer without aldrithiol-4 and incubated with 1 mM of 2,2,2-trifluoroethanethiol (TET) for 1 h at 4 °C. The suspended membranes were pelleted using ultracentrifugation at 200,000 * g for 30 min, resuspended in the same buffer without TET, and incubated with 1 mM theophylline and in-house prepared protease inhibitor solution for 30 min at 4 °C. The protein was extracted by mixing the resuspended membranes 1:1 (v/v) with solubilization buffer (50 mM HEPES pH 7.0, 500 mM NaCl, 0.5% (w/v) n-Dodecyl-β-D-Maltopyranoside (DDM), and 0.05% cholesteryl hemisuccinate (CHS)) for 6 h at 4 °C. The insolubilized material was separated by ultracentrifugation at 200,000 * g for 30 min, and the supernatant was incubated overnight at 4 °C with Co^2+^-charged affinity resin (Talon, Clontech) and 30 mM imidazole.

After overnight incubation, the resin was washed with 20 CV of wash buffer 1 (50 mM HEPES pH 7.0, 10 mM MgCl_2_, 30 mM imidazole, 500 mM NaCl, 8 mM ATP, 0.05% DDM, and 0.005% CHS), and twice with 20 CV each of wash buffer 2 (25 mM HEPES pH 7.0, 250 mM NaCl, 30 mM imidazole, 5% glycerol, 0.05% DDM, 0.005% CHS, and ligand). A_2A_AR[A289C] was eluted with a buffer containing 50 mM HEPES pH 7.0, 250 mM NaCl, 300 mM imidazole, 5% glycerol, 0.05% DDM, 0.005% CHS, and ligand. The eluted protein was exchanged into a buffer (25 mM HEPES pH 7.0, 75 mM NaCl, 0.05% DDM, 0.005% CHS, 100 μM TFA, and ligand) using a PD-10 desalting column (Cytiva) for use in all further experiments. All buffers were prepared with a saturating concentration of ligand.

### Preparation of large unilamellar vesicles with and without A_2A_AR

Phospholipids (POPC or mixtures of POPC and POPS) were dissolved in 2 mL of chloroform. The solution was then evaporated in a rotovap to make a lipid film and then vacuum-dried for 16 h. The lipids were resuspended via repeated vortexing in liposome buffer (25 mM HEPES, pH 7.0, 75 mM NaCl, 100 μM TFA, and ligand) to a final concentration of 1 mM. The lipid mixture was subjected to 9 freeze-thaw cycles to form multilamellar vesicles. Thawing was carried out in a water-bath with a temperature of 10 °C above the highest transition temperature of the lipids used in the mixture. The multilamellar vesicles were then passed through an extruder with a 100 nM polycarbonate filter (Anatrace) 35 times to generate homogenous vesicles.

Vesicles containing A_2A_AR were prepared by first mixing homogenous vesicles with dI H_2_O containing 0.25% DDM and 0.025% CHS for 4 h with gentle rotation at room temperature. The purified receptor was then added to maintain a lipid-to-protein (L/P) molar ratio of 40:1 and allowed to incubate at 4 °C for 4 h. Following this, the sample was then mixed with pre-washed bio-beads (Bio-Rad Laboratories) and incubated for 12-16 hours at 4 °C. The biobeads were removed and the vesicles were passed through an extruder 11 times. These samples were then used for further experiments.

### Dynamic light scattering (DLS) measurements

Lipid vesicles without and with reconstituted A_2A_AR were prepared as described above and diluted one-to-one by volume with liposome buffer (25 mM HEPES, pH 7.0, 75 mM NaCl) to prevent multiple scattering events. DLS experiments were carried out using a Malvern Zetasizer Nano-ZS instrument maintained at 25 °C using cuvettes containing 1.0 mL of vesicles in buffer. The duration of each experiment was 60 seconds. The intensity distributions were calculated with the Zetasizer Software 7.3 (Malvern Panalytical). The polydispersity index of the samples used in this study was maintained at less than 0.1. For each reported data set, three replicates were prepared, and each measurement was performed in triplicate (i.e., 9 measurements in total) to ensure reproducibility.

### Receptor Orientation in Vesicles

The orientation of A_2A_AR[A289C] in LUVs was quantified by adapting a previously described protocol^54^. A_2A_AR[A289C] was purified in aqueous buffer containing DDM/CHS and in complex with ZM241385, as described above, and position C289 was labelled via maleimide chemistry with DY647P1-03 (Dynomics GmbH, Jena, Germany). The purified protein was incubated with 10 molar excess of DY647P1-03 for 3 h at 4 °C in the dark. The reaction was stopped by diluting the sample with 5 X volume of buffer without dye (25 mM HEPES pH 7.0, 75 mM NaCl, 0.05% DDM, 0.005% CHS, and ligand), and the excess dye was removed by buffer exchange through a PD10 column (Cytiva). The labeled A_2A_AR[A289C] sample was reconstituted into vesicles using the above described protocol. The sample fluorescence intensity was monitored using a BMG plate-reader with an excitation wavelength of 640 nm and emission wavelength of 700 nm. 14 mM TCEP (pH 9.0) was added and incubated for 15 min, and the fluorescence intensity was measured again. After 5 min, a solution of 50% Triton X-100 was added to reach a final concentration of 0.05% Triton X-100 in the sample and allowed to equilibrate for 5 min to disrupt the vesicles. After 5 min, the fluorescence intensity was measured again. The orientation of the receptor was calculated from the ratio of the two consecutive quenching steps.

### Electron Microscopy imaging

Liposome samples were prepared with ZM241385 as described above, and the samples were diluted 5X in buffer containing 25 mM HEPES, pH 7.0, 75 mM NaCl. Glow-discharged 400 mesh carbon coated Formvar copper grids (FCF400CU-UB, Electron Microscopy Sciences, Hatfield, PA) were floated onto 5 µl of sample for 5 minutes. The grid was then transferred to a drop of deionized water for 5 s, and then the excess solution was blotted from the grid with filter paper. The sample grid was then floated onto a drop of 1% aqueous uranyl acetate (Mallinckrodt, St. Louis, MO) for 30 s, and then blotted dry. The grids were examined with a FEI Tecnai G2 Spirit Twin TEM (FEI Corp., Hillsboro, OR) and digital images were acquired with a Gatan UltraScan 2k x 2k camera and Digital Micrograph software (Gatan Inc., Pleasanton, CA. The images were processed and analyzed using ImageJ.^67^

### Radioligand binding experiments and data analysis

Saturation binding and competition binding experiments were recorded following previously described protocols.^15,68^ For saturation binding experiments, increasing concentrations of [^3^H]ZM241385 (250 µCi/mmol, Revvity) from 0.1 nM to 20 nM were incubated with 0.1 μg of liposomes containing A_2A_AR[A289C] at 25 °C in buffer containing 25 mM HEPES pH 7.0 and 75 mM NaCl. Specific binding of A_2A_AR[A289C] was determined as the difference in observed binding between samples prepared in the absence and presence of 10 µM ZM241385. All experiments were conducted with three or more replicates. The K_D_ and B_max_ values were determined by fitting the data to a one site binding model in Prism 8 (GraphPad Software, Inc.).

For competition binding experiments, vesicle preparations were incubated with [^3^H]-ZM241385 (2 nM, Revvity) and increasing concentrations of cold ZM241385 or NECA at 25°C for 60 min in buffer containing 25 mM HEPES pH 7.0, 75 mM NaCl. Binding reactions were terminated by filtration through PerkinElmer Easytab-C self-aligned filtermats under reduced pressure using a FilterMat universal harvester (Revvity) and followed by washing twice with 1 mL cold buffer. Radioactivity was measured using a MicroBeta2 microplate scintillation counter (Revvity). All competition binding experiments were conducted with three or more replicates. IC_50_ values were determined using a nonlinear, least-square regression analysis (Prism 8; GraphPad Software, Inc.). The IC_50_ values were converted to K_I_ values using the Cheng−Prusoff equation^69^. Error bars for each measurement were calculated as the standard error of mean (s.e.m) for experiments done in triplicate.

### MAS SSNMR sample preparation

Vesicles containing A_2A_AR[A289C] were pelleted by centrifugation at 200,000 * g for 1 h. The pellet was then packed into a Bruker 1.9 mm rotor by centrifugation at 20,000 * g for 60 min using a series of micropipette tips inserted into the rotor. The total mass for each sample, including lipids, receptor, and aqueous buffer, weighed approximately 15 mg.

### ^19^F Solid-State NMR Spectroscopy

All ^19^F-SSNMR spectra were recorded using a Bruker Avance III HD spectrometer operating at 600 MHz ^1^H nutation frequency using Topspin 3.6.2 and equipped with a Bruker 1.9 mm ^1^H/^19^F/X/Y MAS probe. The flowrate was maintained at 800 liters per hour or higher to maintain temperatures. The 90° ^19^F channel pulse length was 2.33 μs. Two-pulse phase-modulated (TPPM)^70^ decoupling or alternative ^1^H decoupling schemes were applied during the acquisition period, as described in the main text. The heteronuclear decoupling sequence TPPM15 (for 20 kHz and 30 kHz MAS) was used at both high-power (105 kHz) and low power (1/4 MAS) ^1^H decoupling during acquisition. A 30° phase cycling (-15° to 15°) was used for the TPPM sequence for data collection at 40 kHz MAS. Spectra were recorded with a data size of 4k complex points, with an acquisition period of 147 µs, 11k scans, 3.6 µs dwell time, and 2 s recycle delay for a total experimental time of about 6 h per experiment.

An estimation of the sample temperature and calibration of the contribution of frictional heating to the sample temperature was obtained by measuring the ^79^Br chemical shift and spin-lattice relaxation time in a sample of powder KBr at multiple spinning frequencies following previous protocols^61^.

### NMR Data Analysis

All NMR data were processed identically in Topspin 4.0.8 (Bruker Biospin). The ^19^F-NMR data were zero-filled to 32k points and multiplied by an exponential window function with 80 Hz line broadening prior to Fourier transformation. ^19^F-NMR spectra were referenced to the signal from trifluoroacetic acid (TFA) at -75.8 ppm, which was set to 0 ppm. Deconvolution of the ^19^F-NMR data followed previously published procedures and was done with MestreNova version 14.1.1-24571 (Mestrelab Research). Following previously published procedures^14–16^, for each spectrum, the residual difference between the experimental data and sum of the deconvoluted signals was assessed to check the quality of the deconvolution. The relative population of the different A_2A_AR conformational states were calculated as a ratio of integrated area of the each deconvoluted peak to the total integral of all the signals from 6 ppm to 16 ppm.

## ACKNOWLEDGEMENTS

This work was supported by the National Institutes of Health, NIGMS MIRA grant R35GM138291 (M.T.E., B.J., A.P.R.) and by an NIH equipment supplemental grant R35GM138291-03S2. A portion of this work was supported by an NIH award, S10 OD028753, for magnetic resonance instrumentation. The authors also acknowledge the UF ICBR Electron Microscopy Core, RRID:SCR_019146.

## AUTHOR CONTRIBUTIONS

M.T.E. designed the study. B.J. and A.P.R. performed protein production, purification, and vesicle sample preparation. A.P.R. and B.J. recorded and analyzed NMR data with input from M.T.E. M.T.E., B.J. and A.P.R. wrote the manuscript.

## DATA AVAILABILITY STATEMENT

Data used for analysis or generation of figures in this manuscript are available upon request.

## ADDITIONAL INFORMATION

### Supplementary Information

The online version contains supplementary material available at **Correspondence** and requests for materials should be addressed to M.T.E.

