## Supplemental Figures for "The conformational equilibria of a human GPCR compared between lipid vesicles and aqueous solutions by integrative ^19^F-NMR"

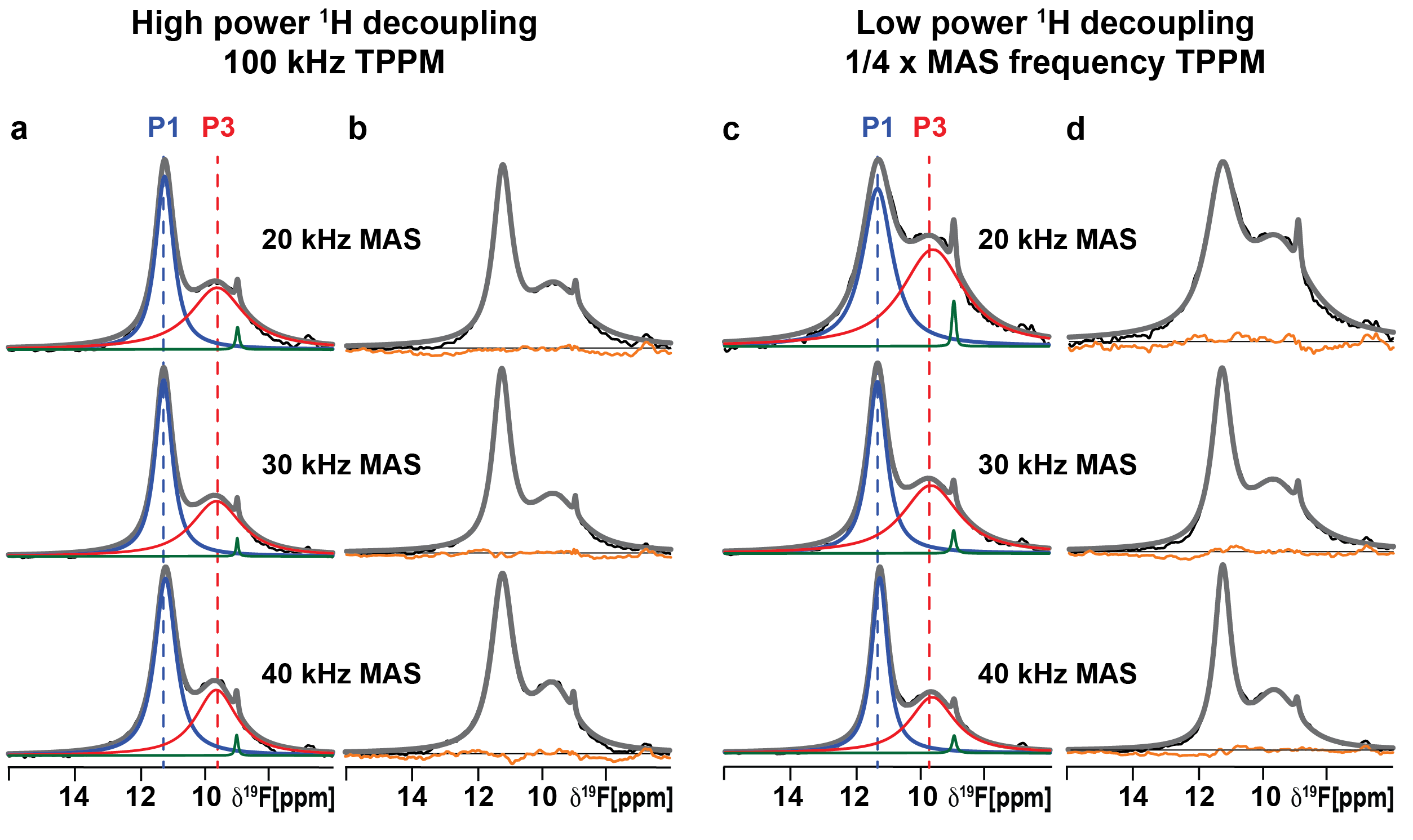

**Figure S1.** **Residual differences between the raw data and summation of the individually fit components from Figure 5.** For each spectrum shown, the experimental data are shown in black, the grey line superimposed on each spectrum is the total sum of the individual deconvolutions, and the orange line is the calculated difference between the grey and black lines. For the spectral components P1 and P3, the same color scheme is used as in Figure 5.

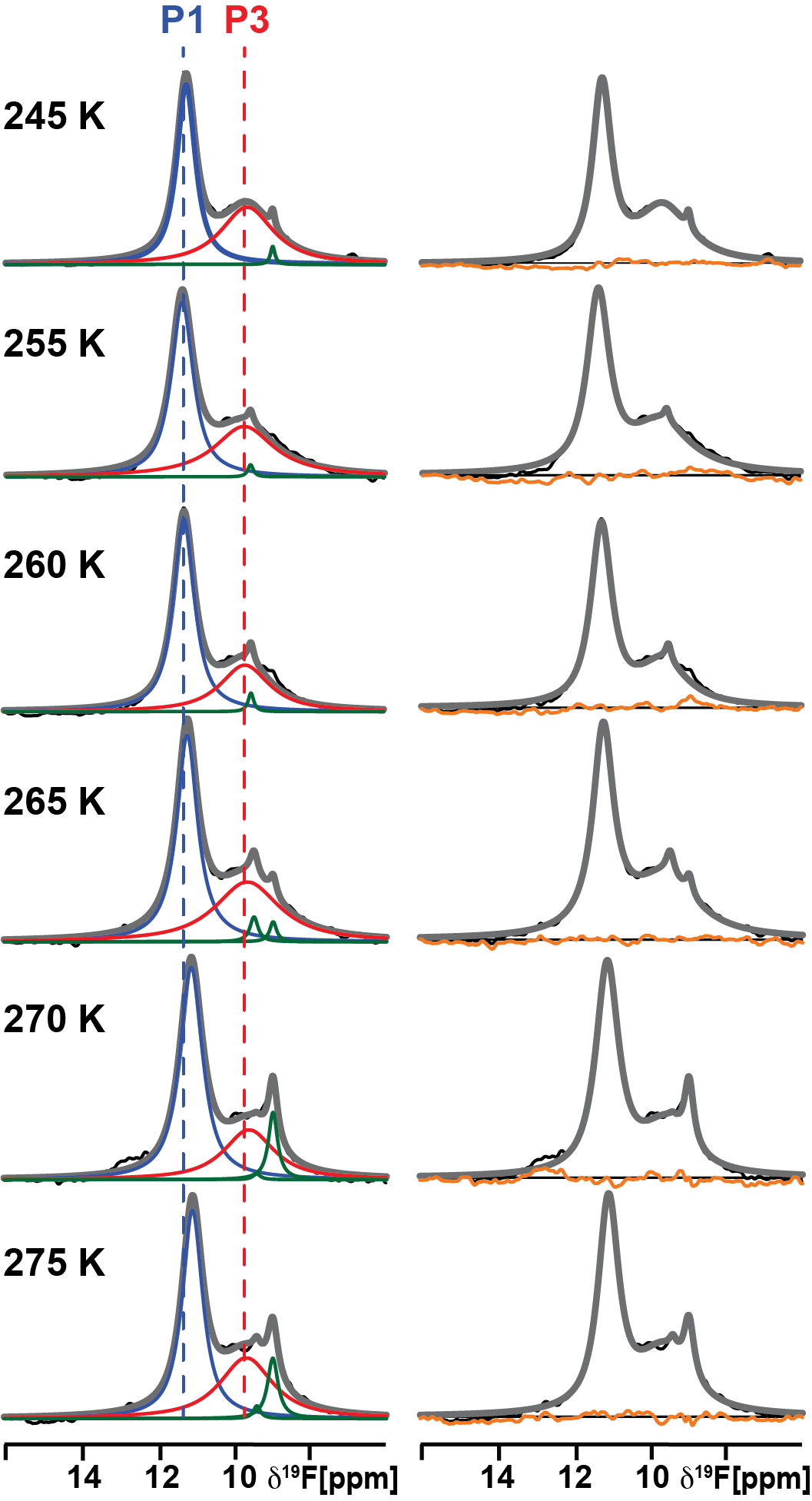

**Figure S2. Variable temperature ^19^F MAS SSNMR spectra of A_2A_AR in complex with the antagonist ZM241385.** The 1-dimensional ^19^F MAS SSNMR spectra are shown for A_2A_AR[A289C] in complex with the antagonist ZM241385 in lipid vesicles containing POPC and POPS (70:30 molar ratio) recorded with 10 kHz ^1^H TPPM decoupling and at an MAS frequency of 40 kHz measured at temperatures ranging from 245 K to 275 K. The reported temperature represents the set point for the temperature controller (see Methods). For each spectrum shown, the experimental data are shown in black, the grey line superimposed on each spectrum is the total sum of the individual deconvolutions, and the orange line is the calculated difference between the grey and black lines. For the spectral components P1 and P3, the same color scheme is used as in Figure 5. The green lines are components from free TET (see text).

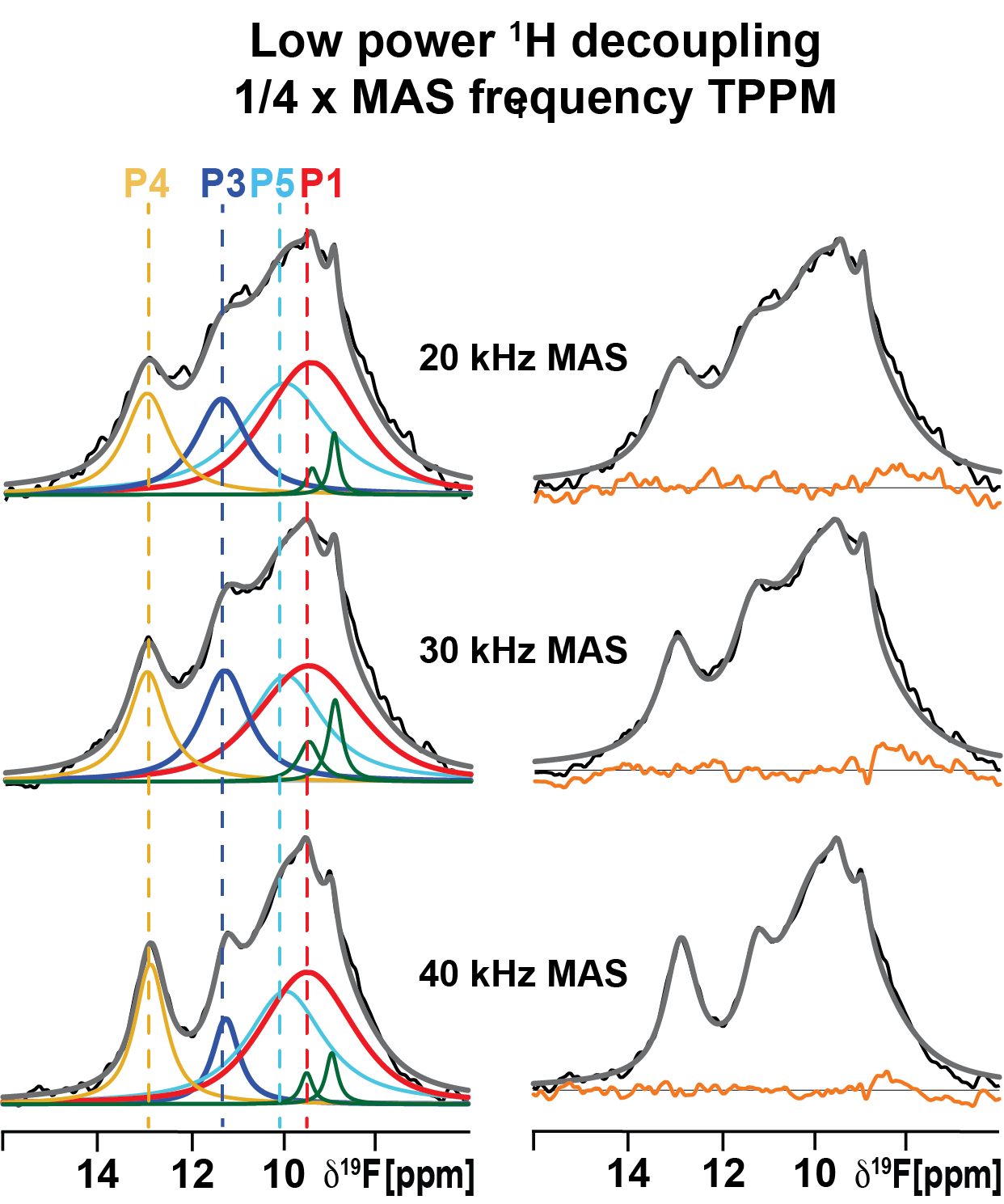

**Figure S3. Residual differences between the raw data and summation of the individually fit components from Figure 6.** For each spectrum shown, the experimental data are shown in black, the grey line superimposed on each spectrum is the total sum of the individual deconvolutions, and the orange line is the calculated difference between the grey and black lines. The same color scheme is used as in Figure 6.

**Table S1: Relative populations and chemical shifts of the A_2A_AR conformational states observed in ^19^F-SSNMR spectra**

The relative populations and the chemical shifts of the A_2A_AR conformational states as observed in the ^19^F-SSNMR spectra are tabulated below. The relative population value reported for each conformation is a ratio of the integrated area of the specific deconvoluted peak to the total integral of all signals from 6 ppm to 16 ppm, excluding signals from free TET.

| Ligand | Spinning  frequencies | Temperature | Decoupling Power | Chemical shifts (in ppm) | | | | | | Relative population | | | |
| --- | --- | --- | --- | --- | --- | --- | --- | --- | --- | --- | --- | --- | --- |
|  |  |  |  | P4 | P3 | P5 | P1 | TET I | TET II | P4 | P3 | P5 | P1 |
| ZM241385 | 20 kHz | 265 K | No decoupling | - | 11.25 | - | 9.59 | 8.93 | - | - | 0.52 | - | 0.48 |
|  |  |  | Low power | - | 11.28 | - | 9.57 | 8.93 | - | - | 0.44 | - | 0.56 |
|  |  |  | High power | - | 11.18 | - | 9.56 | 8.93 | - | - | 0.47 | - | 0.53 |
|  | 30 kHz | 255 K | No decoupling | - | 11.28 | - | 9.56 | 8.93 | - | - | 0.48 | - | 0.52 |
|  |  |  | Low power | - | 11.28 | - | 9.64 | 8.93 | - | - | 0.46 | - | 0.54 |
|  |  |  | High power | - | 11.21 | - | 9.58 | 8.94 | - | - | 0.53 | - | 0.47 |
|  | 40 kHz | 245 K | No decoupling | - | 11.26 | - | 9.66 | 8.96 | - | - | 0.54 | - | 0.46 |
|  |  |  | Low power | - | 11.24 | - | 9.62 | 8.95 | - | - | 0.53 | - | 0.47 |
|  |  |  | High power | - | 11.14 | - | 9.58 | 8.96 | - | - | 0.58 | - | 0.42 |
| NECA | 20 kHz | 265 K | Low power | 12.96 | 11.36 | 10.01 | 9.43 | 8.94 | 9.41 | 0.15 | 0.16 | 0.33 | 0.35 |
|  | 30 kHz | 255 K | Low power | 12.96 | 11.29 | 9.98 | 9.48 | 8.92 | 9.48 | 0.16 | 0.19 | 0.28 | 0.37 |
|  | 40 kHz | 245 K | Low power | 12.83 | 11.21 | 9.92 | 9.45 | 8.93 | 9.47 | 0.17 | 0.09 | 0.34 | 0.40 |

**Table S2: Line widths of the A_2A_AR conformational states observed in ^19^F-SSNMR spectra**

The line widths of A_2A_AR signals observed in ^19^F-SSNMR spectra of various samples are tabulated below. Each value reported in Hz is the full width at half maxima of the ^19^F-SSNMR peak.

| Ligand | Spinning  frequencies | Temperature | Decoupling Power | Linewidth (in Hz) | | | | | |
| --- | --- | --- | --- | --- | --- | --- | --- | --- | --- |
|  |  |  |  | P4 | P3 | P5 | P1 | TET I | TET II |
| ZM241385 | 20 kHz | 265 K | No decoupling | - | 655 | - | 1174 | 89 | - |
|  |  |  | Low power | - | 618 | - | 1293 | 78 | - |
|  |  |  | High power | - | 378 | - | 1131 | 68 | - |
|  | 30 kHz | 255 K | No decoupling | - | 474 | - | 1199 | 69 | - |
|  |  |  | Low power | - | 397 | - | 1175 | 74 | - |
|  |  |  | High power | - | 359 | - | 1034 | 45 | - |
|  | 40 kHz | 245 K | No decoupling | - | 373 | - | 952 | 60 | - |
|  |  |  | Low power | - | 336 | - | 940 | 81 | - |
|  |  |  | High power | - | 441 | - | 857 | 60 | - |
| NECA | 20 kHz | 265 K | Low power | 661 | 750 | 1300 | 1344 | 132 | 168 |
|  | 30 kHz | 255 K | Low power | 557 | 691 | 1080 | 1472 | 200 | 314 |
|  | 40 kHz | 245 K | Low power | 452 | 385 | 1120 | 1306 | 173 | 178 |

**Table S3: Relative populations and chemical shifts of conformational states of A_2A_AR bound to the antagonist ZM241385 observed in variable temperature ^19^F-NMR with low-power TPPM ^1^H decoupling.**

The relative populations and chemical shifts of conformational states of A_2A_AR[A289C] in complex with the antagonist ZM241385 observed in the ^19^F-SSNMR spectra measured with low-power ^1^H TPPM decoupling at different temperatures are tabulated below. The relative population value reported for each conformation is a ratio of the integrated area of the specific deconvoluted peak to the total integral of all signals from 6 ppm to 16 ppm, excluding signals from free TET.

| Ligand | Spinning  frequencies | Temperature | Decoupling Power | Chemical shifts (in ppm) | | | | Relative population | |
| --- | --- | --- | --- | --- | --- | --- | --- | --- | --- |
|  |  |  |  | P3 | P1 | TET I | TET II | P3 | P1 |
| ZM241385 | 40 kHz | 245 K | Low power | 11.24 | 9.62 | 8.95 | - | 0.53 | 0.47 |
|  |  | 255 K |  | 11.33 | 9.70 | - | 9.53 | 0.55 | 0.45 |
|  |  | 260 K |  | 11.26 | 9.66 | - | 9.50 | 0.67 | 0.33 |
|  |  | 265 K |  | 11.21 | 9.64 | 8.96 | 9.46 | 0.52 | 0.48 |
|  |  | 270 K |  | 11.11 | 9.60 | 8.97 | 9.40 | 0.71 | 0.29 |
|  |  | 275 K |  | 11.08 | 9.67 | 8.97 | 9.39 | 0.57 | 0.43 |

**Table S4: Line widths of conformational states of A_2A_AR bound to the antagonist ZM241385 observed in variable temperature ^19^F-NMR with low-power TPPM ^1^H decoupling.**

Line widths of the signals observed in ^19^F-SSNMR spectra of A_2A_AR[A289C] in complex with the antagonist ZM241385 with low-power ^1^H TPPM decoupling at different temperatures are tabulated below. Each value reported in Hz is the full width at half maxima of the ^19^F-SSNMR peak.

| Ligand | Spinning frequency | Temperature | Decoupling Power | Linewidth (in Hz) | | | |
| --- | --- | --- | --- | --- | --- | --- | --- |
|  |  |  |  | P3 | P1 | TET I | TET II |
| ZM241385 | 40 kHz | 245 K | Low power | 336 | 940 | 81 | - |
|  |  | 255 K |  | 396 | 1077 | - | 98 |
|  |  | 260 K |  | 377 | 872 | - | 101 |
|  |  | 265 K |  | 373 | 1133 | 121 | 143 |
|  |  | 270 K |  | 406 | 887 | 163 | 143 |
|  |  | 275 K |  | 367 | 936 | 185 | 127 |
